# Wheat blast caused by *Magnaporthe oryzae* pv. *Triticum* is efficiently controlled by the plant defence inducer isotianil

**DOI:** 10.1101/2020.06.16.154088

**Authors:** Katharina Portz, Florencia Casanova, Angelina Jordine, Stefan Bohnert, Andreas Mehl, Daniela Portz, Ulrich Schaffrath

**Affiliations:** Department of Plant Physiology, RWTH Aachen University, 52056 Aachen, Germany; Bayer AG, Division Crop Science, 40789 Monheim, Germany

**Keywords:** wheat blast, plant defence inducer, spike infection, isotianil

## Abstract

Wheat blast caused by *Magnaporthe oryzae* pv. *Triticum* (*MoT*) is an upcoming threat to wheat cultivation worldwide. The disease crossing over to wheat first gained attention in South America, with increasing interest coming from its more recent appearance in the big wheat growing areas of Asia. The increasing economic relevance of the disease and the lack of genetic resistance in current wheat breading material, besides fungicide resistance already present in fungal pathogen populations, highlighted the need to evaluate the potential of isotianil as an alternative plant protection measure. Isotianil is already registered in Asia for the protection of rice against *M. oryzae* but because the agronomic practices and disease development of blast differ between rice and wheat, the efficacy of isotianil against wheat blast was hard to predict. Testing isotianil in the currently available formulations, applied either as seed treatment or soil drench, resulted in a significant reduction of disease severity. The efficacy was comparably high, on different wheat cultivars and using several fungal isolates with different degrees of virulence. Microscopic analyses revealed that isotianil treatment can prevent invasive growth of the pathogen. No phytotoxicity from isotianil treatment was observed on wheat plants up to the stage of heading. Importantly, isotianil not only protects wheat plants at the seedling stage but also on spikes thereby preventing losses due to this most severe disease syndrome. In summary, the results showed the high potential of isotianil to protect against wheat blast.

## Introduction

Soon after humans began to cultivate plants, devastating plant pathogens emerged, hanging over the heads of farmers like the sword of Damocles. While mankind has striven for a long time to overcome these threats e.g. thanks to the builder of the first green revolution, Norman Borloug, who successfully fought against the wheat stem rust (Rajaram 2011), the challenge remains, as diseases evolve and adapt; this is especially true in the case of novel races with enhanced virulence, such as the so-called warrior race of yellow rust (*Puccinia stiiformis*) or race Ug99 of stem rust (*Puccinia graminis* f.sp. *tritici*, *Pgt*) (Ellis et al. 2014). Additionally, pathogens can re-emerge in areas where they were thought to be eradicated e.g. stem rust (*Pgt*) having an unexpected outbreak in Germany during 2013 (Olivera Firpo et al. 2017) and appearing in the UK after being absent for almost six decades (Saunders et al. 2019). Furthermore, climate change may also add to the increased importance of plant pathogens by favouring the introduction of novel, invasive species into habitats with non-adapted plant communities.

A particular example of the above mentioned scenarios is the occurrence of wheat blast, a disease caused by a specific lineage of the Magnaporthe species complex (Gladieux et al. 2018). This group is best known because of *Magnaporthe oryzae,* a pathogen of rice causing dramatic yield losses of about $66 billion annually, equivalent to the amount needed to feed 60 million people (Pennisi 2010). Wheat blast was first observed in South America in 1985 and spread throughout the whole continent reaching Bolivia in 1996 and Argentina in 2012 (Cruz and Valent 2017). Until recently, the disease was absent from the big wheat growing countries in North America, Europe and Asia but the situation changed dramatically in 2016 and 2017 when it appeared in Bangladesh and India, respectively, most likely via seed imports from South America (Islam et al. 2016). After the disease was reported in Asia, both public and academic attention was raised (Sadat and Choi 2017). An outcome of this situation is the recent debate about the correct naming of the fungus causing wheat blast. While a group headed by Bruce McDonald favors a scenario in which the wheat blast fungus forms a novel species with the name *Pyricularia graminis-tritici* (Castroagudín et al. 2016; Ceresini et al. 2019), a large international consortium led by Barbara Valent, Nick Talbot and Yukio Tosa, all renowned experts in the field of rice blast disease, strongly recommend to treat the wheat blast fungus as a recent tribe (pathotype) of the species *Magnaporthe oryzae* (asexual state: *Pyricularia oryzae*) (Zhang et al. 2016; Cruz and Valent 2017; Valent et al. 2019). For this study we will refer to the pathogen as *M. oryzae* pathotype *Triticum* (*MoT*) (Ceresini et al., 2019; Valent et al., 2019). It is worth noting, that whilst there is an academic discussion regarding nomenclature, there are also far-reaching and very serious practical implications e.g. on international quarantine regulations, which should not be underestimated.

At farmer level, the economic consequence of infection can also be catastrophic; under favourable climatic conditions, the disease can cause up to 100% yield losses not least because grains from infected plants are often deformed or shriveled (Cruz and Valent 2017). The latter happens because the pathogen is not only able to infect leaves but also spikes. Most severely, spike infection may also affect the rachis resulting in obstruction of regular grain filling. A low negative correlation between seedling resistance on leaves and adult plant resistance on spikes was reported (Martínez et al. 2019). As mentioned above, the disease can be transmitted by infected seeds, impacting global quarantine measures. Of course, the best way to avoid accidental transmission of the disease is by preventing infection. Because cultivar resistance against wheat blast was not systematically explored in the past, disease control has generally relied upon application of foliar fungicides. Here, the widespread use of strobilurin chemistry already resulted in fungicide resistance present in pathogen populations e.g. in Brazil (Castroagudin et al. 2015). Moreover the intensive application of fungicides to control other diseases inadvertently selected for *MoT* isolates with resistance against QoIs (e.g. strobilurins) and DMIs (e.g. azoles) (Ceresini et al. 2019). Therefore, some new, alternative approaches are needed to control this disease. In the light of this necessity, we evaluated the efficacy of isotianil, a plant protection agent developed jointly by Bayer CropScience and Sumitomo Chemical Co., Ltd. for the control of rice blast disease and commercially launched during 2010 in Japan and Korea. Although approved only for protection of rice against rice blast or bacterial leaf blight, the disease control spectrum is much broader and covers bacterial and fungal pathogens (Toquin et al. 2012). Because isotianil has no direct antimicrobial effect but instead activates typical plant defence reactions, it has been listed as a plant defence inducer or PDI (Toquin et al. 2012; Bektas and Eulgem 2015; Jeschke 2016). This group of compounds triggers the adaptive plant immune system (Conrath 2009). Here, we report on a study in which the capacity of isotianil as a measure against wheat blast was evaluated. Different application methods were tested for efficacy on both leaves and spikes, on various combinations of wheat cultivar susceptibility and isolate virulence.

## Materials and Methods

### Plant material and fungal isolates

Seeds of wheat plants were germinated on wet filter paper for 24 h and then transferred to standard soil (type ED73, Balster Einheitserdewerk GmbH, Froendenberg, Germany). Seedlings were kept in a growth chamber with 16 h light-(200-250 μmol s-1m-2) and 8 h dark-rhythm at 18 °C and 65 % relative humidity. The cultivar (*cv*.) ‘Little Club’ was obtained from seed stocks in the institute while *cv*. Apogee was received from Eckhard Koch, JKI Darmstadt, Germany. Apogee is a wheat cultivar with a very short life cycle, flowering after 25 days which makes it an ideal choice to study diseases on spikes under lab conditions (Li et al. 2017; Wunderle et al. 2012).

Fungal isolates were placed on dry filter paper at −80°C for long term storage. Isolates for current use were cultivated on oatmeal agar (20 g l^−1^ agar, 2 g l^−1^ yeast extract, 10 g l^−1^ starch, 30 g l^−1^ oat flakes) at 23 °C in the dark. Sporulation was induced by subjecting fungal cultures to a 16h/8 h light/dark regime under black light at 22 °C. The *MoT* isolates Br116.5 and BR32 were obtained from Yukio Tosa, Kobe University, Japan and Didier Tharreau, CIRAD Montpellier, France, respectively. The isolates AR06 and AR33 used in this study were obtained from a collection of *MoT* isolates maintained at CIDEFI (Centro de Investigaciones de Fitopatología) which belongs to the Facultad de Ciencias Agrarias y Forestales - Universidad Nacional de La Plata, Buenos Aires, Argentina.

### Plant inoculation

Preparation of inoculum was according to the method of Delventhal et al. (2014). In brief: conidia were harvested from plates two weeks after sub-cultivation under black light by rinsing the plates with water and filtering through gauze. Concentration of conidia were counted using a Thoma chamber and adjusted to a final concentration of 250,000 conidiospores ml^−1^ in spraying solution (2 g l^−1^ gelatin, 1 ml l^−1^ Tween). For inoculation at the seedling stage, the conidial suspension was sprayed onto leaves and plants, which were then kept in the dark at 24-26°C and 100 % relative humidity for 24 h. Thereafter plants were transferred to a growth chamber, still covered with a plastic hood and kept under the conditions previously described. Spike inoculation was carried out two days after full emergence of the spike. Each spike was separately inoculated with one ml of spore suspension and immediately covered with a pre-wetted plastic bag to maintain high humidity. Plants were placed in the growth chamber, with bags remaining in place, for three days before removal. Preparation of leaf samples for microscopy and assignment of cellular interaction phenotypes to particular categories is described in (Delventhal et al. 2014).

### Treatment with isotianil

Isotianil was applied as either the seed treatment formulation FS 200 or the drench formulation SC 200. For seed treatment, 5 g of wheat seeds were placed in a Falcon tube and covered with 50 μl of a 1:4 dilution of FS 200 in distilled water. This amount of isotianil equates to 50 g a.i. (active ingredient) per dt (decitonnes). Seeds were mixed with the diluted formulation until completely covered. Drench application (13 mg or 26 mg a.i. per pot) was performed by soaking each pot (350 ml soil volume) with 40 ml of isotianil-suspension (57 μl or 114 μl isotianil SC 200 diluted in 40 ml distilled water). For this procedure each pot was placed on an open petri-dish before soaking and kept separately until complete absorbance of the solution. For spray application, an isotianil suspension was prepared by adding 100 μl of isotianil SC 200 to 60 μl distilled water which equals 200 g a.i. per ha (hectare). The solution was sprayed until ‘run-off’ i.e. no more could be retained by the plant surface.

### Quantitative disease scoring

Wheat leaves with clearly visible disease symptoms were laid flat on water agar plates (1% (w/v) agar in distilled water). Pictures of ten leaves per treatment were photographed under indirect light with a camera avoiding light reflections. Quantification of the diseased leaf area was made with the APS software tool Assess 2.0. At first the threshold was roughly set using the Automatic Panel and thereafter adjusted with the Manual Panel to enable a clear differentiation between healthy and diseased leaf areas. The same settings were applied to all samples from each experiment. Values of the diseased leaf area are given as percentages of the total leaf area.

### Quantitative real-time PCR

Total RNA was extracted from secondary wheat leaves (one sample consisting of two leaves, treated as described above) using the citric acid protocol described by Mogga et al. (2016);the material was suspended in 600 μl cell lysis buffer and 200 μl precipitation buffer was added to separate proteins and DNA from the RNA. After that, the RNA was precipitated from the supernatant with isopropanol, washed in 70% ethanol and dissolved in double-distilled water. After digestion of DNA with DNase I (Thermo Fisher Scientific Inc., Germany), cDNA synthesis was performed for each RNA sample (1 μg RNA) using reverse transcription (RevertAid Reverse Transcriptase, Thermo Fisher Scientific Inc., Germany) with HindAnchorT-primer. RT-qPCR was done using iTaq Universal SYBR Green Supermix (Bio-Rad Laboratories Inc. USA), the CFX384 Touch Real-Time PCR Detection System (Bio-Rad Laboratories Inc. USA) and the following gene specific primers: WCI-2_F 5’- ATCACGAGCCAGCTGCAAA-3’, WCI-2_R 5’- GCCTTTTTCGCCTTGACATC-3’ (Sardesai* et al. 2005), EF1α_F 5’-ATGATTCCCACCAAGCCCAT-3’, EF1α_R 5’- ACACCAACAGCCACAGTTTGC-3’ (McGrann et al. 2015). Reactions were heated to 95°C for 3 min, followed by 40 cycles of 95°C for 10 s and 59°C for 30 s. After the final PCR cycle, the reactions were terminated by heating up to 95°C for 10 s and a melting-curve analysis was carried out for each sample. Transcript abundance of the target gene ‘*WHEAT CHEMICALLY INDUCED 2*’ (*WCI-2*) was calculated relative to the transcript abundance of the reference gene ‘elongation factor 1α’ (EF1α) (2^(Ct(reference)-Ct(target))^) (Livak and Schmittgen 2001).

## Results

The wheat blast pathogen *MoT* is able to cause disease symptoms on leaves and ears of infected wheat plants. A causal relationship between sporulation of the fungus on leaves and the later occurrence of ear infections is still a matter of debate and accordingly, it is questioned whether screening for resistance against *MoT* on wheat leaves provides reliable answers for resistance expressed on ears. In response to these unsolved questions, we decided to test the effectiveness of isotianil against wheat blast on both plant organs, i.e. leaves and ears. As known from other pathosystems, the efficacy of isotianil can vary depending upon product placement, therefore seed dressing, soil drench and foliar spray applications were all included.

### Isotianil effectively protects wheat plants at the seedling stage against *MoT*

Seed treatment was made prior to sowing using two different concentrations of isotianil which equals 10 or 50 g a.i. per dt seed, (IST 10 and IST 50, Fig. 1 and Online Resource 1). Plants derived from these seeds and plants from untreated seeds which served as control (UTC) were inoculated with fungal conidia 17 days after sowing, on the second fully expanded leaf. Disease severity was scored one week after inoculation by quantification of the diseased leaf area using digital imaging. Both the area and length of disease symptoms caused by *MoT* on inoculated leaves of isotianil treated plants were significantly reduced compared to untreated control plants (Online Resource 1A). Evaluation of the efficacy of seed treatment with isotianil was carried out over four independent experiments. To enable comparability between replicates, the UTC-value for diseased leaf area was set in each experiment to 100% (Online Resource 1a). A significant reduction of diseased leaf area on isotianil treated plants could be shown in all biological replicates. This reduction was on average 70% and 82% for a treatment with 10 and 50 g a.i. per dt, respectively. Importantly, the degree of protection was equally good, regardless of the disease severity on control plants (UTC) which varied between experiments in a range from 1-40% (Online Resource 1b). The results from quadruple experiments were calculated as relative reduction in diseased leaf area in comparison to UTC and then expressed as mean and standard error in Fig. 1. In a similar way, drench and spray treatments were evaluated in quadruple experiments using different concentrations of isotianil and altering the time span between treatment and inoculation. All results are given for individual experiments (Online Resource 2 and 3) and summarized in Fig. 1. From these results it was obvious that drench treatment performed best in protecting wheat against *MoT*; drenching with solutions of 13 or 26 mg a.i. per pot at five or seven days prior to inoculation resulted in an almost 100% protection rate (Fig. 1). Merely applying 13 mg a.i. per pot at three days prior to inoculation, showed a lower efficacy of 80% reduction of diseased leaf area relative to respective control plants (UTC). This was quite similar to the seed treatment with 50 g a.i. per dt. Noticeably, spray treatments with different concentrations of isotianil at three, five or seven days prior to inoculation gave more variable results and a generally lower degree of protection with 70% reduced infected leaf area being best after treatment with 200 g a.i. per hectare at seven days before inoculation.

**Fig. 1.**
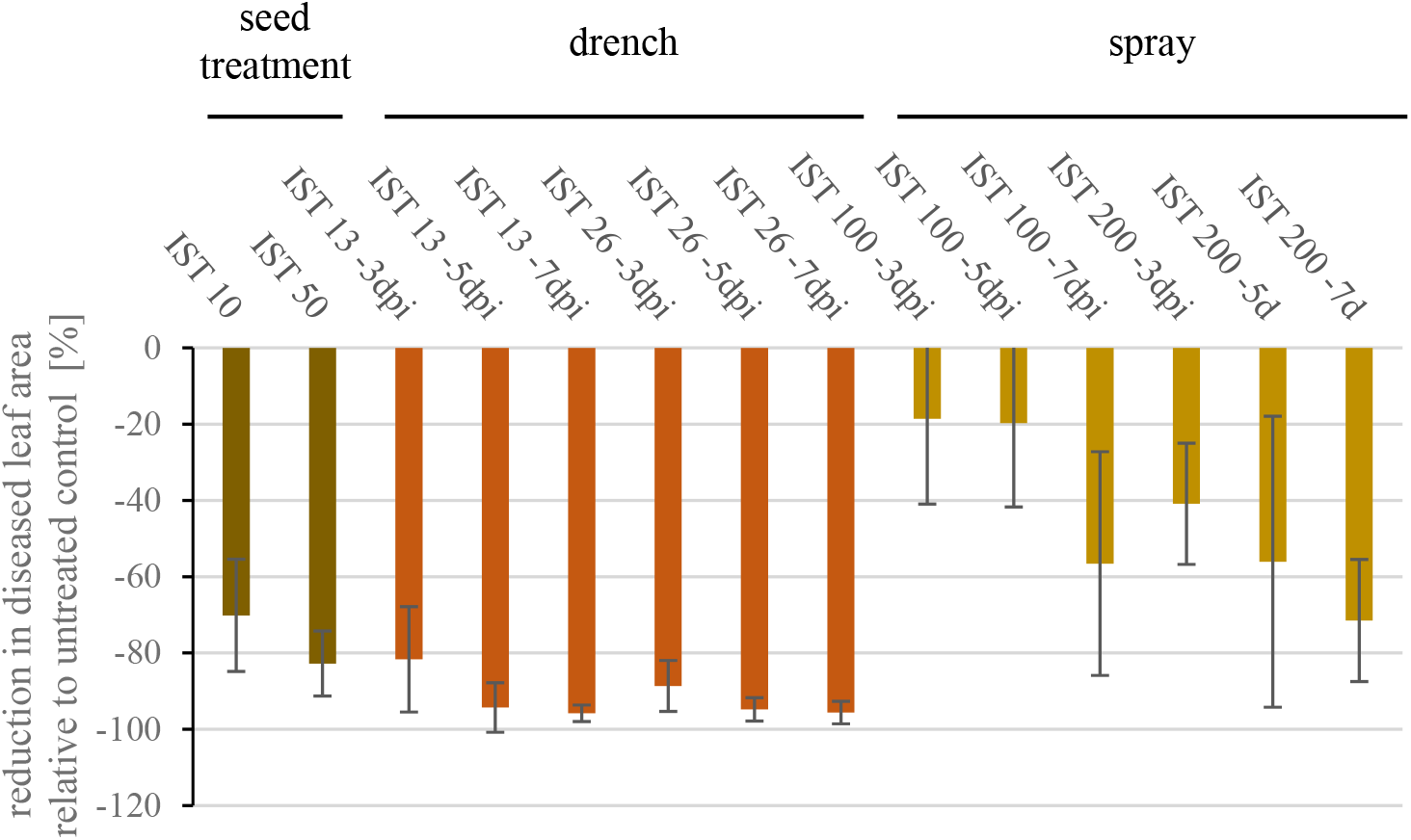
Effectiveness of isotianil in controlling wheat blast using different methods of application. Isotianil was applied to wheat plants *cv*. Little Club either by seed, drench or spray treatment at different concentrations and at varying time points prior to inoculation with the virulent *MoT* isolate BR32. Disease scoring was done on the third leaf one week after inoculation by taking photographs of leaves from 10 plants. Diseased leaf area was determined on photographs and quantitatively evaluated with the software APS Assess 2.0. Values shown are the mean and standard error of four independent biological replicates and represent the reduction of diseased leaf area relative to untreated control plants. Details of individual experiments are given in Online Resources 1-3.

### Aggressiveness of *MoT* isolates did not influence protection by isotianil

Experiments shown above were obtained with the wheat cultivar, ‘Little Club’ and *MoT* isolate BR32. In these tests, although the disease severity on untreated plants varied between biological replicates, this did not influence the efficacy of isotianil treatments (Online Resources 1-3). To broaden the base for this observation the set-up was extended by including an additional wheat cultivar, i.e. Apogee, and three additional isolates of *MoT* (Online Resource 4). Macroscopic evaluation of disease severity revealed the highest aggressiveness for the Argentinian isolates AR06 and AR33 on both wheat cultivars. In contrast, isolates BR32 and BR116.5 showed a lower infection rate on *cv.* Little Club compared with *cv.* Apogee (Online Resource 4). Because drench application was shown to be most effective in previous experiments (Fig. 1), this technique was used to apply isotianil, at a concentration of 13 mg a.i. per pot, five days prior to inoculation of the second leaf. In all cases, treated plants were clearly less severely infected following artificial inoculation *MoT* (all isolates on both wheat cultivars). Quantitative assessment of this experiment was carried out in an analogous way to the results already presented in Fig. 1. This evaluation substantiated the macroscopic observations by confirming that both wheat cultivars were similarly highly susceptible to the Argentinian *MoT* isolates AR06 and AR33, with leaves showing 40-45% diseased area (Fig. 2, a). By contrast, leaves of wheat cultivar Apogee were less severely affected by the Brazilian isolates BR32 and BR116.5 at only 5 and 10% of diseased leaf area, respectively, with *cv*. Little Club being even less susceptible at approximately 5% diseased leaf area from either isolate (Fig. 2, A). Remarkably, the degree of protection after drench application of isotianil was best on both wheat cultivars, when attacked by the most aggressive isolates AR06 and AR33 (Fig. 2, b). Nonetheless, even on plants which are only slightly infected in this particular experiment, i.e. Apogee and Little Club after inoculation with BR32, a 60% reduction in diseased leaf was recorded.

**Fig. 2.**
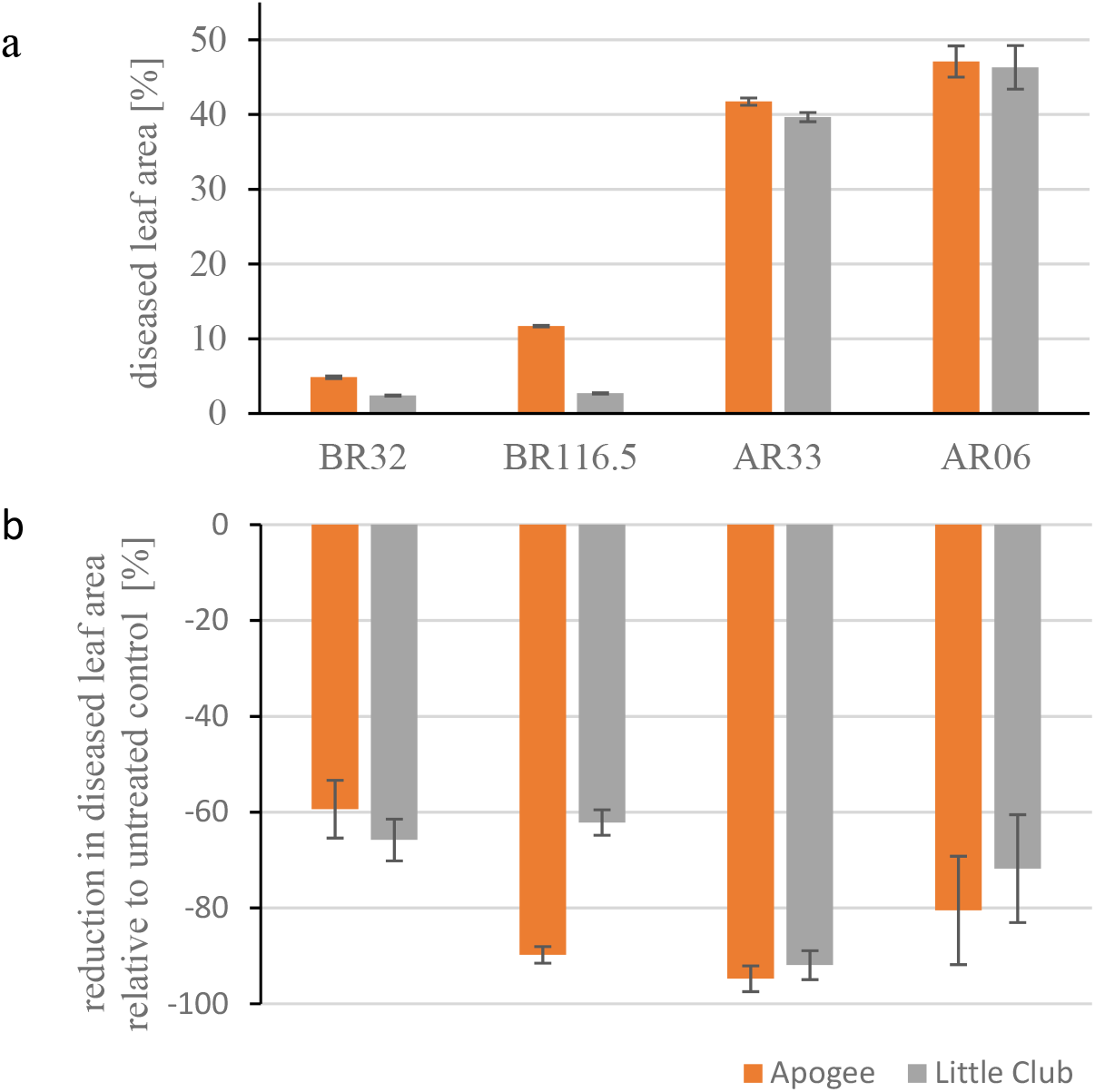
Degree of susceptibility of wheat cultivars against different isolates of *MoT*. The two wheat cultivars Apogee and Little club were tested for their response towards different isolates of the wheat blast fungus. The isolates originated either from Brazil (BR32 and BR116.5) or Argentina (AR33 and AR06). Wheat plants were grown until full emergence of the second leaf and then ten plants were treated with isotianil at 13 mg a.i./pot by drench, whereas the other ten plants remained untreated. After additional five days, all plants were inoculated with conidia of isolates as indicated. Disease scoring was made on the third leaf one week after inoculation by taking photographs of ten leaves. Diseased leaf areas were determined on photographs and quantitatively evaluated with the software APS Assess 2.0. Values shown are the mean and standard error. a) Comparison of susceptibility for both wheat cultivars to *MoT* isolates. b) Reduction of diseased leaf area on plants treated with isotianil relative to untreated plants. Pictures of disease symptoms on treated and untreated plants are shown in Online Resource 4.

Extensive use of fungicides may cause shifts in pathogen populations towards isolates showing fungicide resistance which eventually is correlated with a fitness penalty (Hawkins and Fraaije 2018). It is well known that *MoT* populations in Brazil display resistance to strobilurin fungicides (so-called quinone-outside inhibitors, QoI) by non-synonymous nucleotide substitutions in the cytochrome b gene that prevents fungicide binding. While the G143A mutation, occuring at a frequency of 36%, provides complete fungicide resistance, the F129L mutation confers moderate resistance (Castroagudin et al. 2015). To check for the possibility that the Brazilian isolates BR32 and BR116.5 contain these mutations and therefore are less aggressive on their host, as similarly reported for *M. oryzae* on perennial ryegrass (Ma and Uddin 2009), a PCR-based analysis was performed (Online Resource 5). The results conclusively showed that neither the G143A nor the F129L mutations were present in isolates used in this study.

### Microscopic analysis revealed that isotianil prevents invasive growth of *MoT*

To determine the mode of action of isotianil on pathogen colonisation, a histological analysis was performed. As reported in previous studies, plant-pathogen interaction sites, i.e. plant cells attacked by the fungus, were inspected and grouped into four different categories which are associated with plant defence or successful pathogen development (Online resource 6) (Delventhal et al. 2014). All microscopic samples were firstly screened under bright field microscopy on located appressoria and then the switch to epi-fluorescent light allowed the determination of auto-fluorescence of the plant cell at the infection site. The deposition of auto-fluorescent material beneath an appressorium can be correlated with a halt of fungal invasion (Jarosch et al. 2005; Jarosch et al. 2003). In contrast, deposition of auto-fluorescent material associated with collapsed mesophyll cells is a sign of successful pathogen progress from epidermal into mesophyll tissue and is presumably associated with transition from biotrophic to necrotrophic life-style. Quantitative assessment of these categories revealed that at 72 and 96 hpi the frequency of infection sites grouped into the fourth category (collapsed mesophyll cells) increased from 5 to 15 % in plants not treated with isotianil (Fig. 3, UTC) which is in accordance with successful infection of *MoT* isolate BR32 as shown in Fig 2. By contrast, the frequency of interaction sites grouped into this category were around 1-2 % at 72 and 96 hpi for plants treated with isotianil. This reveals that the pathogen was unable to progress from an initially attacked epidermal cell into the mesophyll and consequently pathogen invasion was stopped.

**Fig. 3.**
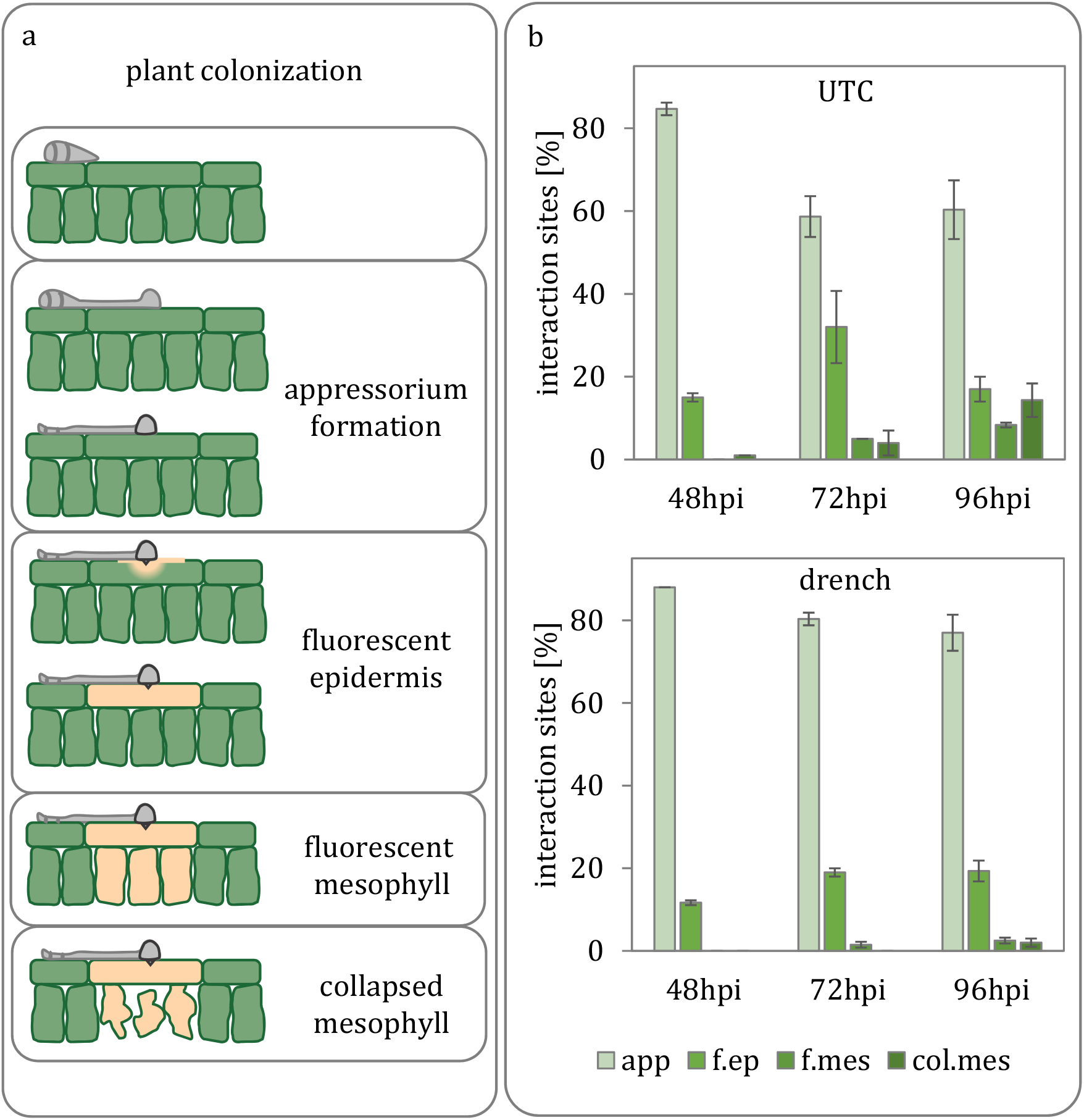
Quantitative microscopy of wheat plants treated with isotianil and inoculated with *MoT*. The primary leaves of seven-day-old wheat seedlings (cv. Little Club) were drenched with isotianil at 13 mg a.i. /pot. Five days after treatment, secondary leaves were inoculated with the virulent *MoT* isolate BR32. Inoculated leaves were harvested at 48 hpi, 72 hpi and 96 hpi and analysed by brightfield and epi-fluorescence microscopy after bleaching. (a) Schematic presentation of the fungal infection process and categories formed in association with plant responses. (b) The diagram shows the frequency of these categories per leaf for plants not treated with isotianil (UTC) or plants drenched with the compound (drench) harvested at different time points after inoculation. Values shown represent the mean ± standard deviation of three leaves consisting of 100 evaluated interaction sites per leaf. Results shown are from a single experiment. hpi, hours post inoculation; app, appressorium; col.mes, collapsed mesophyll; f.ep, fluorescent epidermis; f.mes, fluorescent mesophyll.

### Drench treatment with isotianil at stage of flag leaf development reduces spike infection

The efficacy of isotianil in protecting wheat against *MoT* at the seedling stage has been presented above. However, the pathogen is more aggressive in its infection of the spikes (ears). Experiments were therefore also performed to evaluate the efficacy of seed, drench and spray application in this respect. All experiments with spike infections were carried out with *cv*. Apogee because of the exceptionally short time needed for this cultivar to reach ‘heading’ (Strugala et al. 2015). While seed treatment with isotianil did not show any degree of protection, spray and especially drench application at the stage of fully emerged flag leaves, gave remarkable reductions in the number of spikelets showing bleaching due to *MoT* infection (Fig. 4). The highest degree of protection was achieved by drenching. Inspection of seeds after maturation substantiated this observation (Online Resource 7). According to their size and shape, seeds were categorized either as healthy or shriveled. Control plants neither treated with isotianil nor inoculated with *MoT* had on average 30 healthy seeds per spike. After inoculation the proportion of healthy seeds decreased to 48% and 15% in two different experiments while the average number of seeds per spike remained the same (Online Resource 7a and 7b). Consistent with the macroscopic scoring of infected spikes, drench treatment at the flag leaf stage turned out to provide the highest degree of protection with 79% of healthy seeds per spike (Online Resource 7a). Even more efficient, however, was a combined treatment with two applications of isotianil at first by seed treatment followed by a drench application at the stage of flag leaf expansion. In this case 90% of healthy seeds could be recovered from inoculated spikes.

**Fig. 4.**
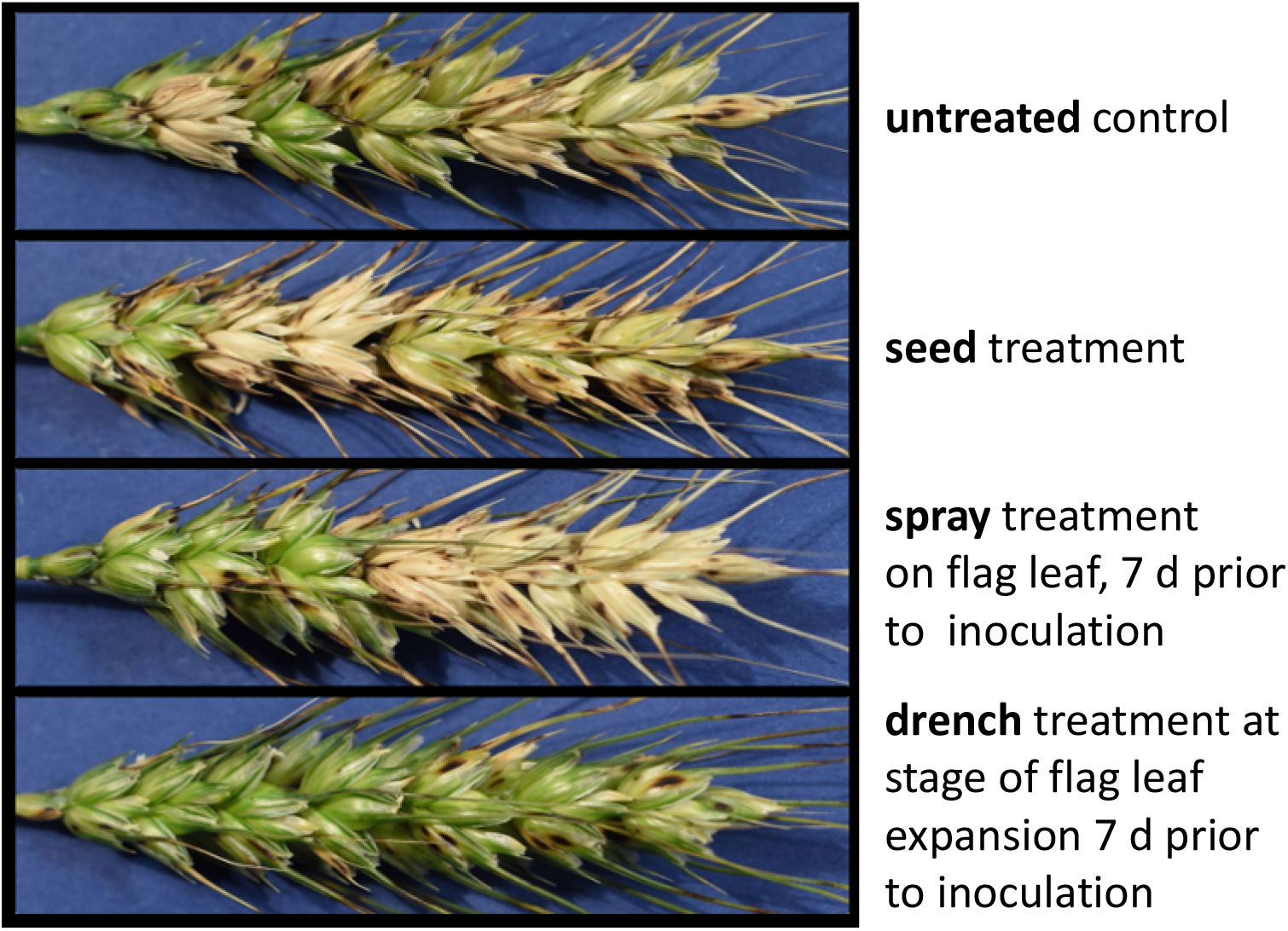
Spike infection with the wheat blast fungus *cv.* Apogee. The effectiveness of isotianil in controlling wheat blast on spikes were evaluated in response to three different types of treatment, i.e. seed, drench and spray application, respectively. While seed coating (50 g a.i./ dt) was done prior to sowing, drenching (200 g a.i./ha) and spraying (26 mg a.i./pot) took place on the fully emerged flag leaves. One week after these treatments spikes had developed and were inoculated with the virulent isolate AR 06. Pictures of spikes were taken 17 days after inoculation.

### Isotianil treatment triggers expression of the wheat lipoxygenase gene *WCI-2*

Plant defence inducers have been intensively studied as protective measure against plant diseases. In the early days, progress in our understanding of the underlying molecular processes was made by correlating transcript abundances with the onset of resistance. At that time, a number of new genes in wheat were described as being induced after treatment with Benzothiadiazole (BTH) and named ‘*WHEAT CHEMICALLY INDUCED*’ (*WCI*) genes (Gorlach et al. 1996). Here, we tested whether the wheat gene *WCI-2*, encoding a lipoxygenase, also responds to isotianil treatment. Therefore, wheat plants were treated with isotianil by drenching at the stage of fully expanded primary leaves (around seven days after sowing). Then quantitative real-time (qRT)-PCR was used to monitor *WCI-2* transcript abundance in the newly emerged second leaves. This revealed a more than ten times increase in transcript abundance at the fourth day after treatment (Fig. 5a). Next, treatment with isotianil was combined with pathogen inoculation. In this assay, plants were first treated with isotianil on the first emerged leaf and then inoculated with a compatible *MoT* isolate after emergence of the second leaf (five days after isotianil treatment). RNA was extracted from the inoculated leaf and qRT-PCR analysis again confirmed a significant accumulation of *WCI-2* transcripts in response to isotianil. However, pathogen inoculation alone did not lead to an increase of *WCI*-*2* transcripts and combining isotianil treatment with inoculation also showed no enhancement of the isotianil-dependent increase (Fig. 5b).

**Fig. 5.**
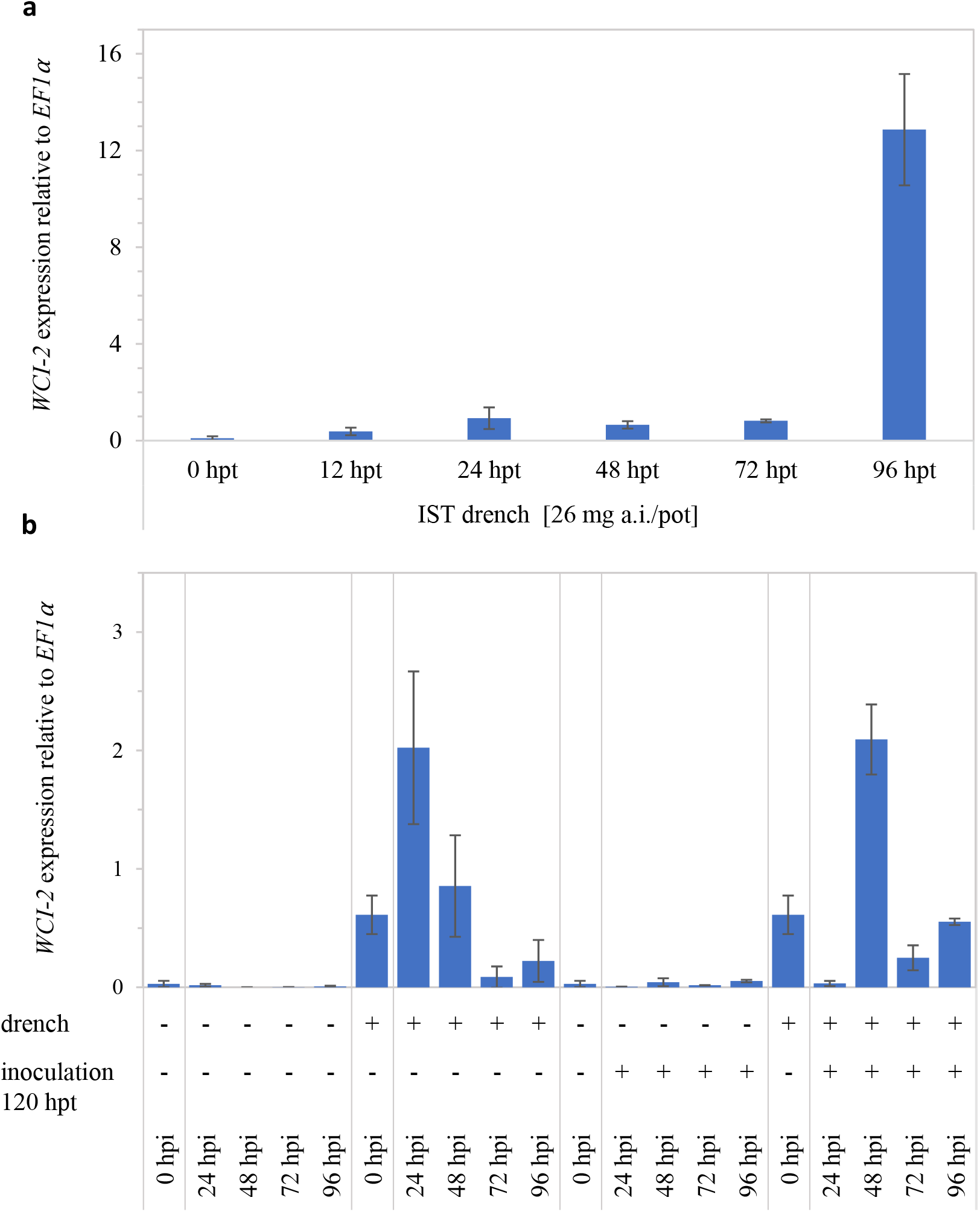
Relative transcript abundance of *WCI-2* after isotianil treatment. Plants of wheat *cv.* Little Club were treated with isotianil seven days after sowing at the stage of primary leaf expansion. a) Treatment was done by drenching each pot with 26 mg a.i. The second leaf from nine individual plants was harvested at time points indicated and used for RNA isolation. b) Plants were drenched with isotianil at 13 mg a.i./pot and five days after treatment (120 hpt) all plants were inoculated on the second leaf with the compatible *MoT* isolate BR32. RNA was isolated from these secondary leaves at different time points after inoculation. RNA from (a) and (b) was transcribed to cDNA and used for quantification of *WCI-2* transcript abundance by RT-qPCR. The expression of *WCI-2* was calculated relative to EF1⍺. Values shown represent the mean ± standard deviation of three samples consisting of two leaves each. Results shown are from a single experiment. hpt, hours post treatment; hpi, hours post inoculation

## Discussion

Wheat blast emerged as novel threat to wheat cultivation and has the potential to tremendously impact global food security. One key question in the debate of how to fight against this disease, is whether our extensive knowledge on blast disease on rice, caused by *Magnaporthe oryzae*, can be used as blueprint for the investigation of wheat blast. Here we present a study on the use of isotianil, a well-known plant protection against rice blast, to control wheat blast. Our results demonstrated that isotianil is similarly effective in wheat as it is in rice.

Isotianil was launched in 2010 in Japan and Korea as a protection against rice blast and bacterial leaf blight (Toquin et al. 2012). Due to the specific requirements of rice cultivation, as a submerged cropping system, the product was optimized either for seed coating or drench application in seed boxes. Accordingly, these two application methods were first evaluated in wheat and were found to provide a highly significant level of protection against wheat blast at the seedling stage (Fig. 1, Online Resource 1 and 2). Doses of 10 mg a.i. per dt seeds and 13 mg a.i. per pot, were shown to be sufficient for seed and drench application, respectively, to gain a 70 to 95% reduction in diseased leaf area (Fig. 1). These doses are of agronomical relevance and similar to those used in rice cultivation (Toquin et al. 2012). Because in wheat cropping systems fungicides are mostly applied by spraying, we additionally evaluated this method of treatment. Spray application, however, turned out to be less effective compared to the aforementioned treatments (Fig 1 and Online Resource 3). This could either mean that the active ingredient is not sufficiently taken up by wheat leaves, potentially due to an inappropriate formulation for the leaf type, or that isotianil is not properly transported within the plant, i.e. first by basipetal and then an acropetal transport mechanism. Nonetheless, results after seed and drench application confirmed the efficacy of isotianil against *MoT* in wheat. Remarkably this holds true even at different disease pressures as exemplified by 1–55% and 6-70% diseased leaf area in the untreated control of experiments with seed and drench treatment, respectively (Online Resource 1 and 2). To further substantiate this observation, we tested the protection provided by drenching with isotianil against *MoT* isolates of different origins. Even against the most aggressive isolates, AR33 and AR06, the reduction in disease severity was similarly high as compared with the less aggressive isolates, BR32 and BR116.5 (Fig. 2, Online Resource 4). These results contradict the argument that synthetic plant defence inducers, such as isotianil, are only effective at low disease pressure. Using quantitative microscopy, it could be demonstrated that the treatment with isotianil led to an inhibition of invasive growth of the pathogen, i.e. colonisation was stopped during transition from epidermal to mesophyll tissue (Fig. 3). To further rule out the possibility that the *MoT* isolates used in this study are less vital, possibly due to the presence of a mutation conferring resistance to strobilurin fungicides (Hawkins and Fraaije 2018), a PCR-based approach was undertaken. Using specific primers for amplification of gene fragments spanning the respective regions, it was shown that neither the G143A nor the F129L amino acid swap were present in any of the isolates (Online Resource 5). These mutations are already quite common in *MoT* populations in Brazil and this already limit the use of classical fungicides and highlight the necessity for alternatives such as isotianil (Castroagudin et al. 2015).

The most severe and even most apparent indication for wheat blast disease in fields are spike infections which, in extreme cases, can lead to complete bleaching of a spike (Cruz and Valent 2017). Because seedling resistance on leaves and adult plant resistance on spikes are not strictly correlated (Martínez et al. 2019), an investigation of both plant stages and organs is necessary. Under lab conditions, experiments with wheat spikes are difficult to conduct because of the long time needed between sowing and flowering and the growing space needed for large wheat plants at the adult stage. In this respect, a particularly useful experimental tool is the spring wheat cultivar ‘Apogee’ which has an exceptionally short life cycle and flowers 25 days after planting, without vernalization (Li et al. 2017). First it was confirmed that *cv.* Apogee was similarly protected by isotianil, against wheat blast at the seedling stage, as was demonstrated for *cv*. Little Club (Fig. 2, Online Resourse 4). In wheat cropping a final fungicide treatment is routinely applied at the flag leaf stage. Therefore, *cv.* Apogee plants were treated at this stage by drench and spray application with isotianil and compared to plants emerged from seeds coated with isotianil. The evaluation of disease symptoms on spikes around three weeks after inoculation revealed that drench application was the most effective treatment (Fig. 4). Monitoring the amount of shriveled seeds per spike gave further support to this observation (Online Resource 7a) linking reduced spike infections to increased yield.

Interestingly, the combination of seed treatment followed by drench application most efficiently reduced the proportion of malformed seeds (Online Resource 7b). This observation might pave the way for the use of isotianil in wheat cropping because it provides a long period of protection from newly emerged seedling to mature plant prior to harvest. It was shown that transcripts of the wheat gene *WCI-2* strongly accumulate after isotianil treatment (Fig. 5). This suggests that isotianil functions as a defence inducer in wheat, as previously shown for BTH (Gorlach et al. 1996). Additionally, it was observed that other diseases of wheat, i.e. powdery mildew, could also be controlled by isotianil (Online Resource 8) which confirmed previously published data (Toquin et al. 2012).

In summary, the presented data has shown isotianil as an interesting new tool for the management of wheat blast disease.

## Acknowledgments

The authors are thankful to Prof. Analia Perello from University La Plata, Argentina for helpful discussions. Dr. Richard Meredith from Bayer AG is kindly acknowledged for proof-reading of the manuscript and helpful discussion.

## Author contribution

KP, FC and AJ performed experiments and evaluated results. DP and AM were involved in the experimental design and helped in finalizing the manuscript. US designed the study, accompanied the experiments, drafted and finalized the manuscript.

