## Supplemental Figures for "Wheat blast caused by *Magnaporthe oryzae* pv. *Triticum* is efficiently controlled by the plant defence inducer isotianil"

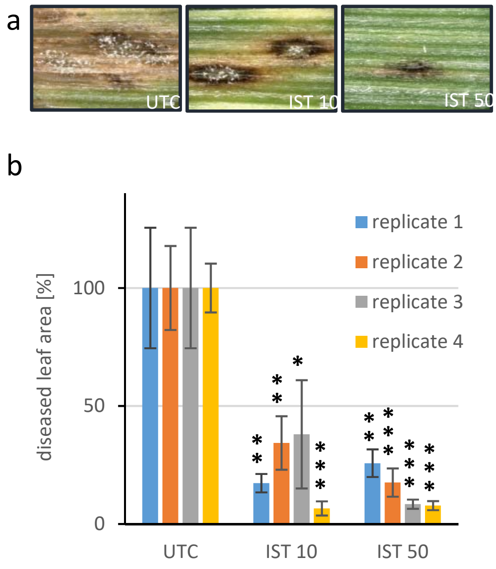


**Online Resource 1. Effectiveness of seed treatment with isotianil in controlling wheat blast.** Isotianil was applied to seeds of wheat plants *cv.* Little Club at 10 or 50 g a.i./dt seeds prior to sowing. Plants were inoculated with the virulent *MoT* isloate BR32 17 days after sowing. a) Representative pictures of individual infection sites are shown for plants not treated with isotianil (UTC, untreated control) and plants treated with 10 or 50 g a.i./dt (IST 10 or IST 50). b) Diseased leaf areas were quantitatively scored on third leaves by taking photographs from 10 plants and evaluation with the software APS Assess 2.0. Values shown are the mean and standard error of four independent biological replicates. In each experiment the value for untreated control plants was set to 100% which equals an absolute value of 15%, 43%, 1% and 43% diseased leaf area in replicate one to four, respectively. For each experiment significant differences between the control and treatments are marked with asterisks as follows: * = p <= 0,05; ** = p <= 0,01; *** = p <= 0,001.

abspeichern als ESM_1.pdf

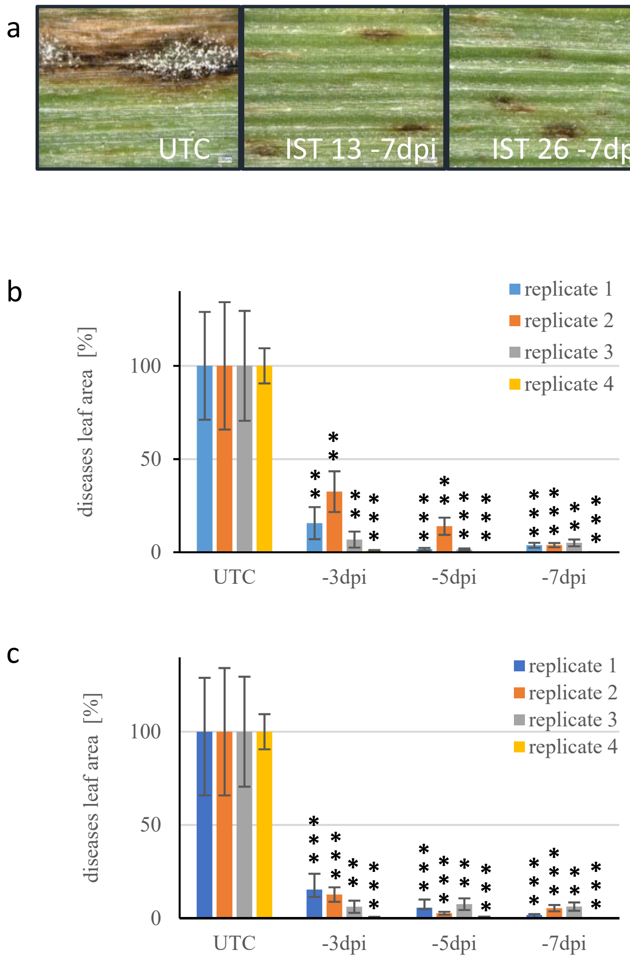


**Online Resource 2. Effectiveness of drench application of isotianil in controlling wheat blast.** Isotianil was applied to wheat plants *cv.* Little Club at three, five or seven days prior to inoculation by drenching with either 13 or 26 mg a.i./pot. Plants were inoculated with the virulent *MoT* isolate BR32 seven days after sowing. a) Representative pictures of individual infection sites are shown for untreated control plants (UTC,) and plants treated with 13 or 26 mg a.i./pot seven days before inoculation (IST 13 -7 dpi or IST 26 -7 dpi). b) Diseased leaf area was quantitatively scored on third leaves of plant treated with 13 mg a.i./pot at three, five or seven days prior to inoculation (dpi) and by taking photographs from 10 plants which were evaluated with the software APS Assess 2.0. Values shown are the mean and standard error from these nine leaves. Results from four independent biological replicates are presented. In each experiment the value for untreated (UTC) was set to 100% which equals an absolute value of 6%, 22%, 1% and 55% diseased leaf area in experiments one to four, respectively. For each experiment significant differences between the control and treatments are marked with asterisks as follows: * = p <= 0,05; ** = p <= 0,01; *** = p <= 0,001. c) Depiction of data obtained for wheat plants treated with isotianil at 26 mg a.i./pot. Evaluation similar to b.

abspeichern als ESM_2.pdf

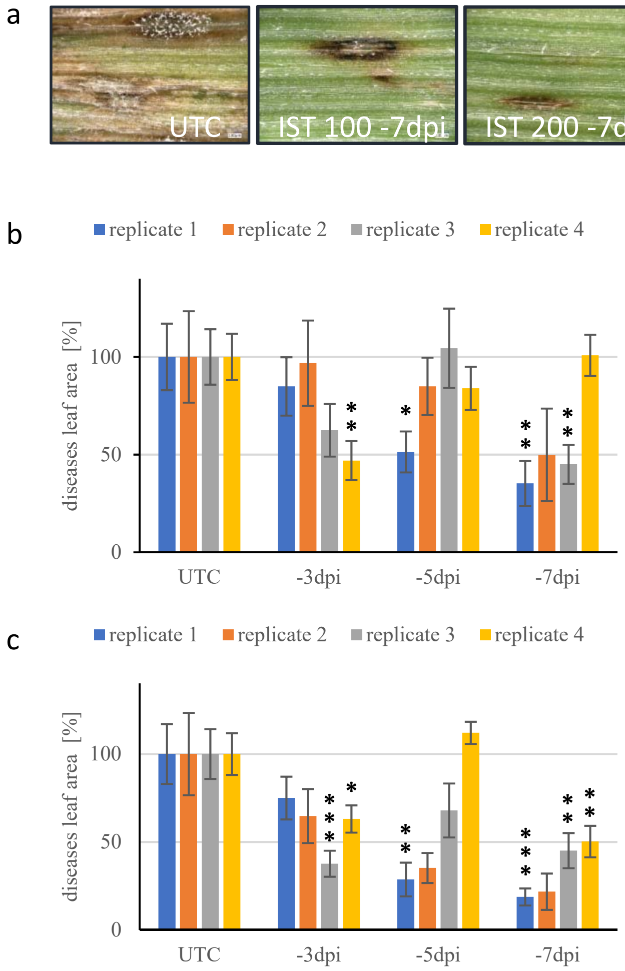


**Online Resource 3. Effectiveness of spray application of isotianil in controlling wheat blast.** Isotianil was applied to wheat plants *cv.* Little Club at three, five or seven days prior to inoculation by spraying a solution with either 100 or 200 g a.i./ha. Plants were inoculated with the virulent *MoT* isolate BR32 seven days after sowing. a) Representative pictures of individual infection sites are shown for plants not treated with isotianil (UTC, untreated control) and plants treated with 100 or 200 g a.i./ha seven days prior to inoculation (IST 100 -7 dpi or IST 200 -7 dpi). b) Diseased leaf areas were quantitatively scored on third leaves of plant treated with 100 g a.i./ha at three, five or seven days prior to inoculation (dpi) and by taking photographs from 10 plants which were evaluated with the software APS Assess 2.0. Values shown are the mean and standard error of four independent biological replicates. In each experiment the value for plants not treated with isotianil (untreated control, UTC) were set to 100% which equals an absolute value of 6%, 31%, 2% and 70% diseased leaf area in experiments one to four, respectively. For each experiment significant differences between the control and treatments are marked with asterisks as follows: * = p <= 0,05; ** = p <= 0,01; *** = p <= 0,001. c) Depiction of data obtained for wheat plants treated with isotianil at 200 g a.i./ha. Evaluation similar to b.

abspeichern als ESM_3.pdf

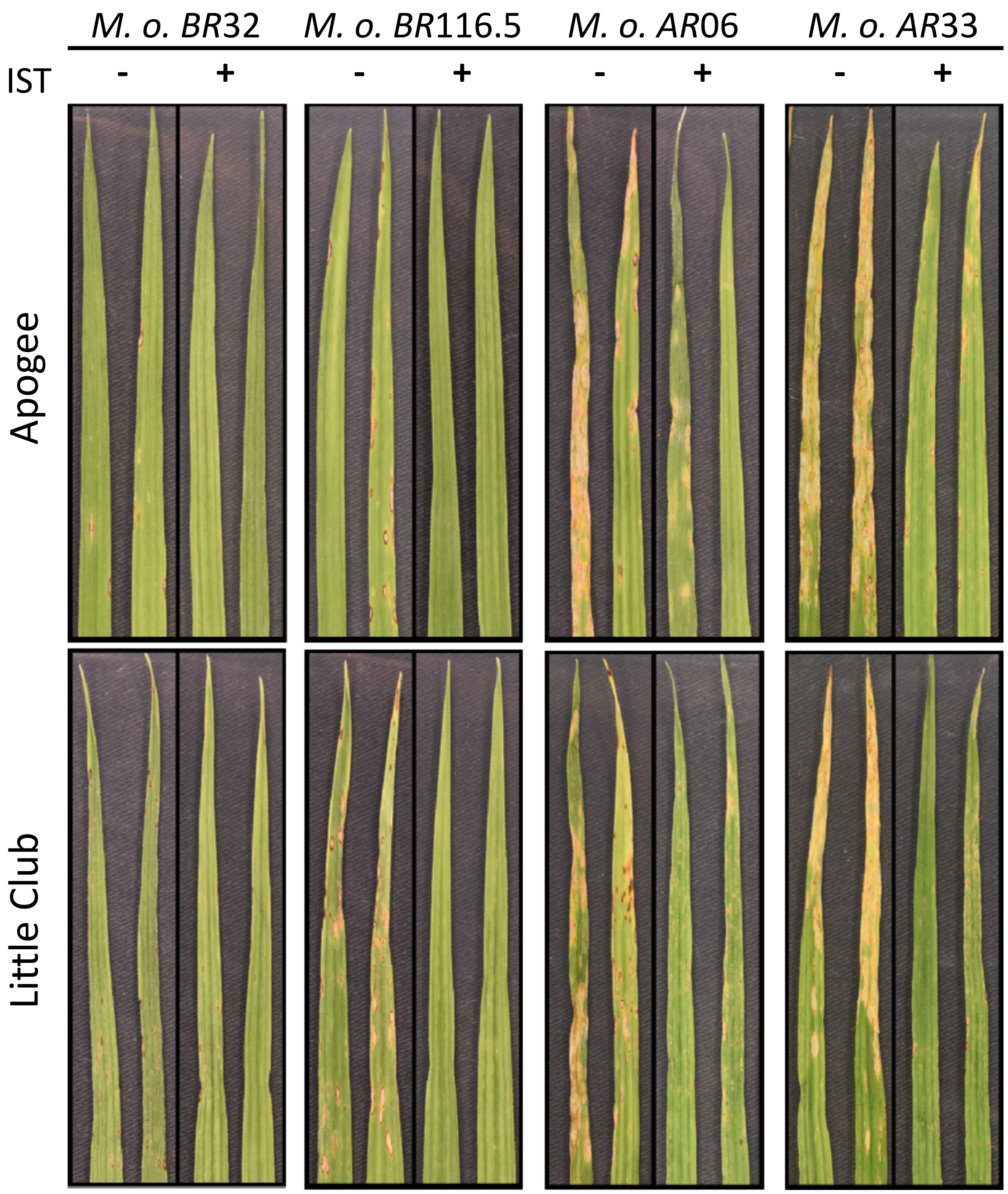


**Online Resource 4. Effectiveness of drench application of isotianil in controlling wheat blast on different wheat cultivars and in response to four wheat blast isolates with different degree of aggressiveness.** Isotianil was applied to plants after full emergence of the second leaf by drench at a concentration of 13 mg a.i./pot and marked with (+) while untreated control plants are indicted by (-). Seven days after treatment plants were inoculated with suspensions of conidia from isolates as indicated. Representative pictures were taken from diseased plants seven days after inoculation.

a) analysis of G143A mutation (GGT to GCT)

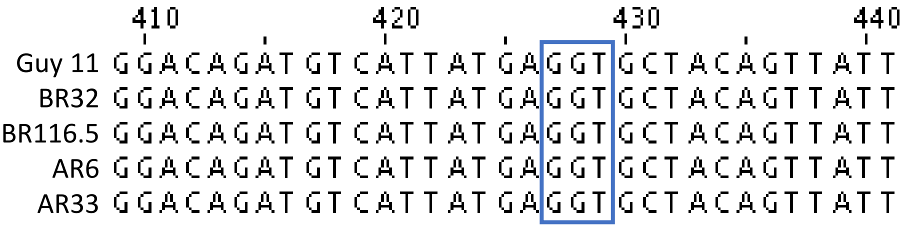


b) analysis of F129L mutation (TTC to TTA)

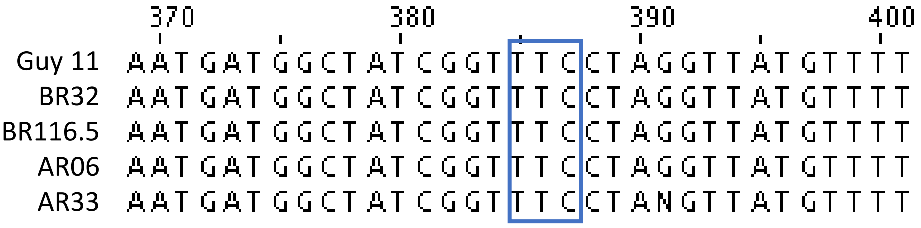


**Online Resource 5. Sequence analysis of *M. oryzae CYTB* revealed no mutations associated with QoI resistance.** A fragment of the cytochrome B gene (*CYTB)* was amplified from genomic DNA by PCR using the primers PgCtyb-F1 (5‘-AGTCCTAGTGTAATGGAAGC) and PgCytb-R1 (5‘ ATCTTCAACGTGTTTAGCACC) and conditions as published in Castroagudin et al., Phytopathology 105:284-294. The PCR products were separated by agarose gel electrophoresis, purified and sequenced. Sequences shown correspond to the region of the gene, boxed in blue, where mutations are known to confer resistance to fungicides from the QoIs group (quinone-outside inhibitors). a) G143A mutation (G-to-C transition at position 428) and b) F129L mutation (C-to-A transversion at position 387). Sequences of the four isolates used in this study were aligned with the Guy 11 *CYTB* sequence obtained from the database using Jalview version 2.10.3b.

Castroagudín, V. L., Ceresini, P. C., de Oliveira, S. C., Reges, J. T. A., Maciel, J. L. N., Bonato, A. L. V., Dorigan, A. F., and McDonald, B. A. 2015. Resistance to QoI fungicides is widespread in Brazilian populations of the wheat blast pathogen *Magnaporthe oryzae*. Phytopathology 105:284-294.

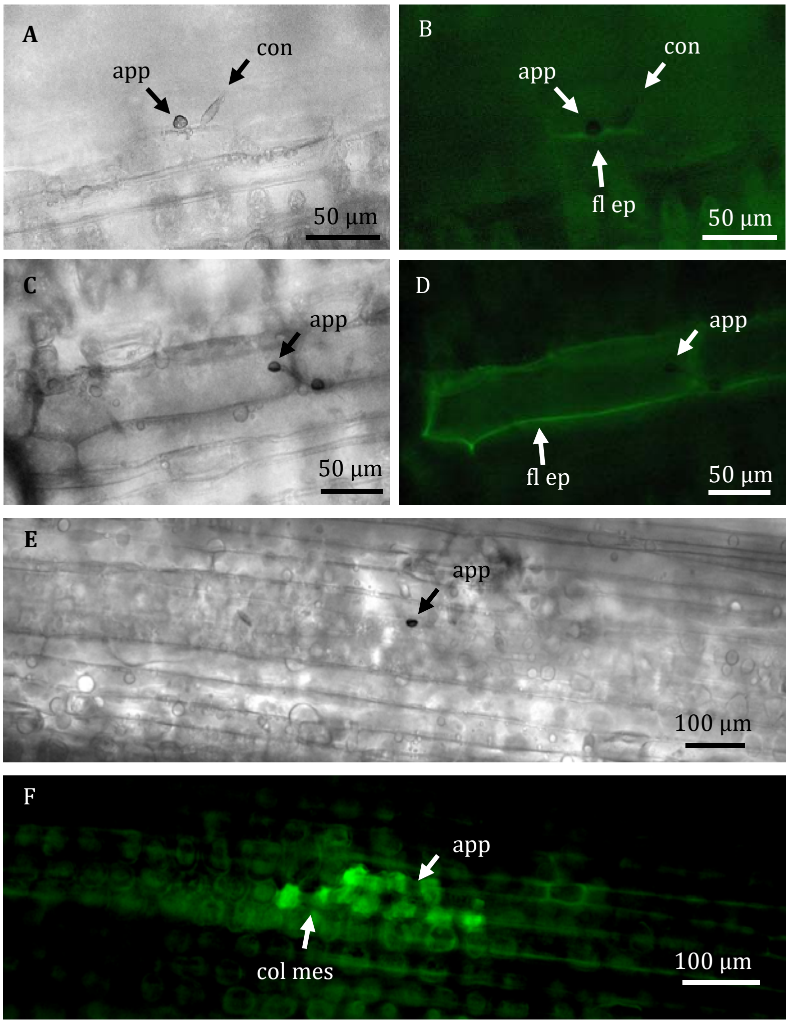


**Online Resource 6. Infection sites of *MoT* on wheat plants.** Secondary leaves of wheat plants (*cv.* Little Club) inoculated with *MoT* isolate BR32 were harvested at different time points. After extraction of leaf pigments samples were evaluated by bright field (A, C and E) and epi-fluorescence microscopy (B, D, and F). Pictures shown represent different categories of plant responses expressed by cells associated with fungal appressoria: (A, B) faint fluorescence of epidermal cell beneath appressorium; (C, D) pronounced fluorescence of cell walls of an epidermal cell beneath appressorium; (E, F) fluorescence of collapsed mesophyll cells beneath an epidermal cell attacked by a fungal appressorium. app, appressorium; col mes, collapsed mesophyll; con, conidium; fl ep, fluorescent epidermal cell

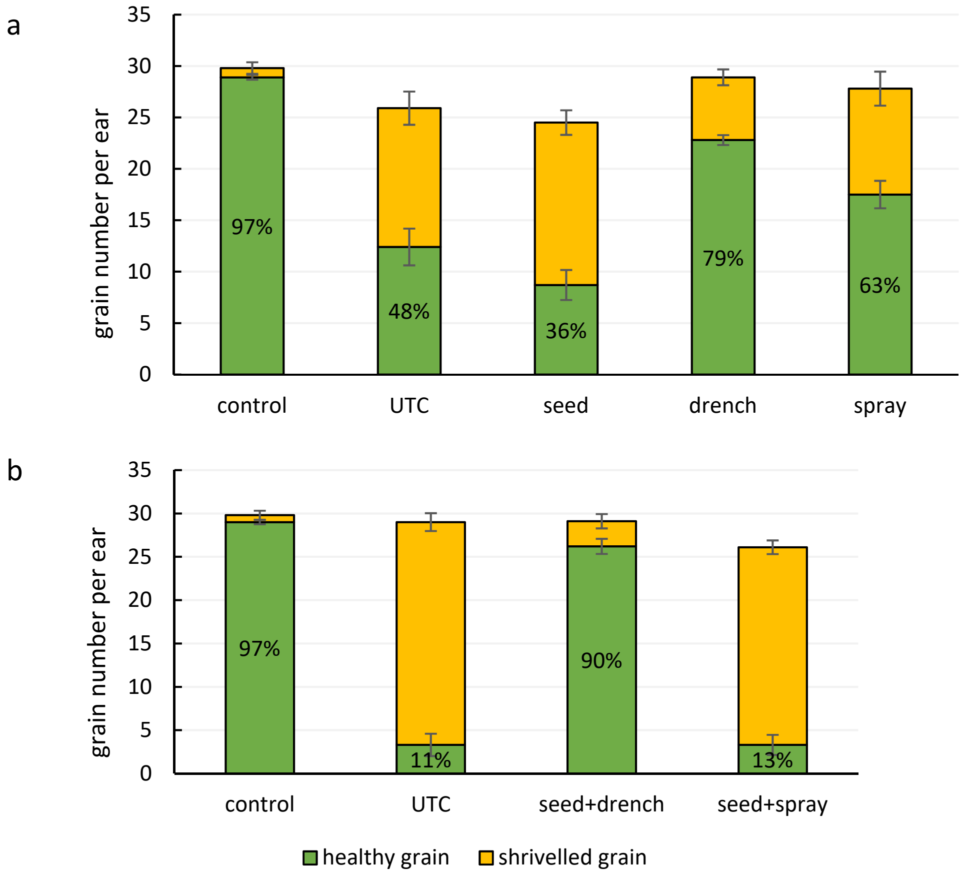


**Online Resource 7. Efficacy of different isotianil treatments on grain yield.** Apogee seeds and plants were treated by three different methods with isotianil. a) Seeds were treated with 50 g a.i./dt, plants at the flag leave stage were sprayed with 200 g a.i./ha or drenched with 26 mg a.i./pot. When spikes were fully emerged (approx. seven days later) plant were inoculated with *MoT* BR32. b) Combined treatments were done as follow: seeds were first coated with 50 g a.i./dt and, after plants reached the flag leave stage, flag leaves were sprayed with 200 g a.i./ha or drenched with 26 mg a.i./pot. For quantitative analysis seeds were harvested and categorized into two groups, shriveled and healthy. Values are the mean and standard error of the seeds from 10 spikes for every treatment method.

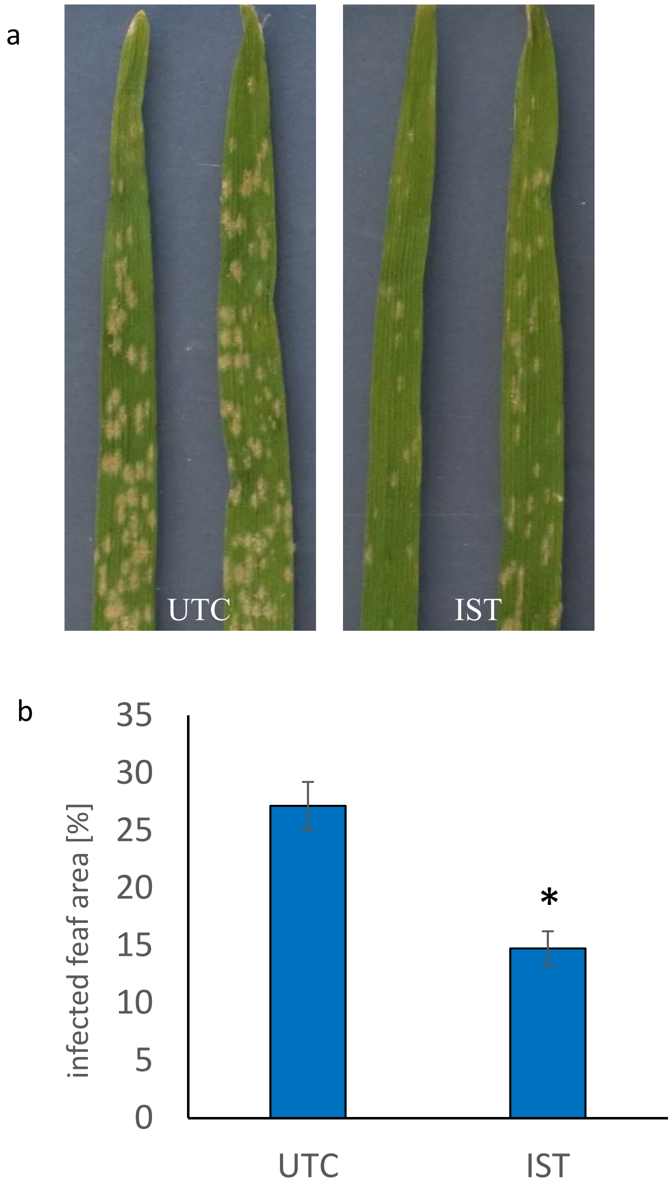


**Online Resource 8.** Effectiveness of drench application of isotianil in controlling powdery mildew on wheat. Wheat plants *cv.* Little Club were treated with isotianil by drenching at 13 mg a.i./pot at the primary leaf stage. Seven days after treatment, plants were inoculated with *Blumeria graminis* f.sp. *tritici* and another seven days later plants were scored for disease symptoms. a) Representative pictures of untreated control plants (UTC) and plants treated with isotianil (IST). b) Diseased leaf area was quantitatively determined using pictures of ten leaves and evaluation with the software APS Assess 2.0. Values shown are the mean and standard error of a single experiment (asterisk, P = <0,001).
